# Paclitaxel sensitizes TRAIL (tumor necrosis factor-related apoptosis-inducing ligand)-resistant breast cancer cells towards TRAIL-mediated apoptosis

**DOI:** 10.64898/2026.03.18.712553

**Authors:** Nirajan Ghosal, Divisha Biswas, Dwaipayan Chaudhuri, Manisha Sarkar, Kalyan Giri, Ranjana Pal

## Abstract

**Background:** The ability of TRAIL to specifically induce apoptosis in cancer cells makes it a promising candidate to be an effective chemotherapeutic drug. But resistance to TRAIL treatment is a major obstacle. Finding combinatorial therapies that make resistant tumors more susceptible to TRAIL is an effective preclinical approach. In this work, we investigated the possibility that pre-treatment of paclitaxel may promote apoptosis in TRAIL-resistant breast cancer cells.

**Methods:** *In silico* analysis was done to investigate the binding affinity between TRAIL receptors (DR5 and DCR2) and paclitaxel via docking and MD simulation. To check whether any non-lethal dose of paclitaxel can modulate the expression of TRAIL receptors, qPCR was done in paclitaxel treated breast cancer cells. Next, paclitaxel was pre-administered to TRAIL-resistant MCF7 and MDA-MB-453 human breast cancer cells followed by rhTRAIL treatment. Cell viability and survival was evaluated using the MTT assay and colony formation assay, respectively. Immunoblot for caspase-3 was performed to study apoptosis. The expression level changes of DR5 and DCR2 were analyzed post-treatment using qPCR and immunoblot assay.

**Results:** *In silico* analysis showed that paclitaxel can bind with higher stability to DCR2 in comparison to DR5 thereby changing the preference of TRAIL molecules towards DR5. Next, in cell line experiments we observed that administering a non-lethal dose of paclitaxel to MDA-MB-231 and MCF7 breast cancer cells resulted in no significant cell death but led to an increase in DR5 and a decrease in DCR2 expression at both the transcript and protein levels. Furthermore, in TRAIL-resistant MCF7 and MDA-MB-453 cells, pre-treatment with paclitaxel followed by rhTRAIL administration induced significant cell death due to paclitaxel induced increase in DR5 as well as decrease in DCR2 expression at both the transcript and protein levels. Moreover, long term survival of MDA-MB-453 cells was significantly lower when pretreated with paclitaxel and exposed to rhTRAIL compared to control, paclitaxel alone or rhTRAIL alone group.

**Conclusion:** Thus, our study uncovers a novel therapeutic strategy to overcome TRAIL resistance underscoring the clinical potential of using a non-lethal dose of paclitaxel to modulate TRAIL receptor dynamics. Future research should be aimed at exploring the potentiality of using paclitaxel-based combinatorial approaches in crafting effective TRAIL therapies.

## Introduction

The worldwide impact of breast cancer is on a significant rise assessed in terms of both incidence and mortality [1]. In the year 2022 itself, more than 2.3 million cases of breast cancer were diagnosed and more than 0.67 million deaths were recorded in females on a global scale. In the same year, age-standardized incidence and mortality rates in Asia were estimated to be 34.3 and 10.5 per 100000, respectively, with 0.95 million new cases and 0.3 million deaths reported. In India alone, breast cancer is responsible for more cancer-related deaths in women in comparison to other cancers [1]. With such high numbers in hand, developing a robust cure associated with the disease or even a standardized treatment regimen is currently both imperative and challenging. On the subject of anti-cancer therapy, a member of the TNF ligand superfamily, Tumor Necrosis Factor Related Apoptosis Inducing Ligand (TRAIL) has become an attractive option for chemotherapeutic medication because it can cause cell death in malignant cells while effectively safeguarding normal cells and eluding systemic as well as lethal in vivo toxicities [2]. There are several TRAIL-based drugs under clinical trial, encompassing anti-TRAIL receptor antibodies, recombinant TRAIL molecules, and gene therapy agents. Recombinant TRAIL (rhApo2L) or the use of TRAIL-death receptor agonists (mapatumumab, apomab, lexatumumab) are the two primary treatment modalities. However, only a small number of medications have advanced to phase II clinical trials, and these methods appear to have limited therapeutic value [3]. Despite these promising attributes, TRAIL has not yet gained widespread adoption in clinical practices due to the problem of TRAIL resistance. TRAIL forms homotrimers and binds to two death receptors, TRAIL-R1 (DR4) and TRAIL-R2 (DR5), as well as two decoy receptors, TRAIL-R3 (DCR1) and TRAIL-R4 (DCR2). On TRAIL binding, the death receptors can initiate the apoptotic pathway with the help of a cytoplasmic region called the death domain (DD). Receptor oligomerization on the surface of cell membrane is followed by the formation of death inducing silencing complex (DISC) which in turn causes activation of caspases, eventually leading to induction of apoptosis. TRAIL resistance or inhibition against TRAIL-induced apoptosis can be explained by the role played by the decoy receptors. DCR1 is a membrane anchored protein and lacks the aforementioned death domain while DCR2 possesses a truncated, non-functional death domain. Therefore the decoy receptors can only bind TRAIL and sequesters it away thus impeding apoptosis [4]. In this light, DCR2 has been implicated in TRAIL resistance not only due to its capability of TRAIL quenching but also by triggering anti-apoptotic genes [5, 6], inducing pro-survival molecular pathways, and promoting cellular proliferation [7–10] thus protecting the malignant cells against TRAIL-mediated apoptosis. Consequently, recent research has focused on developing strategies to overcome this resistance in clinical settings. For instance, the combination of TRAIL and paclitaxel has demonstrated synergistic anti-glioblastoma (GBM) activity *in vivo*, with minimal toxicity to normal tissues [11]. Similarly, co-delivery of the TRAIL gene enhances the antitumor effects of paclitaxel against GBM cells both *in vitro* and *in vivo* [12]. Combining TRAIL with death receptor agonists (e.g., mapatumumab) [13] or natural compounds (e.g., resveratrol) [14] offers a multifaceted approach to sensitize cancer cells, improve therapeutic outcomes, and reduce resistance.

Among the several chemotherapeutic drugs, paclitaxel, also known as taxol, is a potent antimitotic and anticancer drug that has been used for the treatment of breast cancer since 1994 [15]. Initially derived from the Pacific Yew tree (*Taxus brevifolia*), paclitaxel is a plant alkaloid that belongs to the taxane chemical group [16]. It is recognized as a crucial antineoplastic drug, widely utilized in the clinical treatment of various other cancers, including those of the bladder, lungs, head and neck, ovaries and esophagus [17]. As an antimitotic chemotherapeutic agent, paclitaxel stabilizes microtubules by interfering with the dynamic balance between tubulin dimers and microtubules, resulting in cell cycle arrest at the G2/M phase. It is also capable of inducing apoptosis in human cells through multiple pathways including modulation of the anti-apoptotic proteins (Bcl-X_L_), pro-apoptotic proteins (Bax, Bak), mitotic spindle checkpoint-associated apoptotic signal transduction pathways, abnormal activation of cyclin-dependent kinases, JNK/SAPK signaling pathway, transcriptional regulation of apoptosis regulating genes, etc [18]. However, the therapeutic potential of paclitaxel is limited due to multiple dose dependent side effects like coma, acute encephalopathy, neutropenia, granulocytopenia, generalized myalgia, neurotoxicity, cardiotoxicity, gastrointestinal toxicity and even death [19]. According to clinical research, treatment protocols incorporating paclitaxel alongside other anti-cancer medications have shown a 74% improvement in overall survival rates for patients diagnosed with breast cancer [20]. *In-vitro* treatment regimens involving TRAIL and paclitaxel combination treatment resulted in a significant enhancement of apoptosis in prostate cancer [21], ovarian cancer [22], non-small cell lung cancer [23], renal cancer [24], and breast cancer [25]. TRAIL and paclitaxel co-treatment-induced apoptosis occurs through elevated expression of TRAIL receptors (DR4 and DR5) in prostate cancer cells [21]. Apart from this, the combination of TRAIL gene therapy with paclitaxel enhances both anti-tumor and anti-metastasis outcomes in metastatic lung cancer [24]. Also, sequential treatment of paclitaxel followed by TRAIL leads to synergistic cell death in TRAIL-resistant cell lines [26]. Administration of TRAIL-related monoclonal antibody TRA-8 with paclitaxel leads to significant tumor regression in a mouse model of human breast cancer, whereas paclitaxel alone does not do so [27]. Therefore, treatment strategies that involves co-therapeutic approach of TRAIL along with paclitaxel can be promising in overcoming TRAIL resistance. In this study, we have evaluated the potential of paclitaxel as a co-therapeutic agent along with TRAIL to reverse TRAIL resistance in breast cancer.

## Methods

### Structure preparation and Molecular Docking and MD simulation

The complex structure of DCR2 and TRAIL has not been elucidated till date although complex of DR5 and TRAIL is available (1DU3.pdb). Based on structural homology between DCR2 and DR5 receptors, complex structure was built using molecular replacement method in PyMol taking 1DU3.pdb as the template. The paclitaxel molecule was docked to DCR2 and DR5 targeting the interfacial residues of both the complexes using AutoDock Vina and complexes constituting TRAIL, either of the receptors and the paclitaxel molecule was built [28]. Docking was performed using an exhaustiveness of 32 in order to consider most of the rotational structural isomers of the paclitaxel molecule so as to obtain the most stable conformation of the ligand molecule in the protein binding pocket. Four systems were prepared which constitutes TRAIL-DCR2 complex, TRAIL-DR5 complex, TRAIL-DCR2-paclitaxel complex and TRAIL-DR5-paclitaxel complex. The molecular complexes consisting of the protein and ligands were simulated by utilizing the CHARMM36 force field in the GROMACS 2021.5 suite [29]. System preparation was achieved by using the CHARMM-GUI server (Hsu et al., 2017). Solvation of the complexes was done by utilizing the TIP3P water model in a cubic simulation box. For neutralizing the solvated system, sufficient amounts of Na^+^ and Cl^-^ were given, at a concentration of 0.15 M, and the pH was set at 7.4, which is the physiological pH. Applying the algorithm for the steepest descent gradient with a highest value of 50,000 steps, energy minimization of the neutralized system was done. After NVT equilibration was done for 500 ps at 310 K, the system that was now energy-minimized, was exposed to 500 ps of NPT equilibration at 310 K and a pressure of 1 atm. During equilibration, a 1 kcal mol^-1^ [^-2^ force constraint was implemented on the backbone atoms of the protein. Next, the NPT equilibrated systems were exposed to restraint-free 50 ns production runs having 2 fs time steps. The Smooth Particle-Mesh-Ewald (PME) method was implemented for computing the long-range electrostatic interactions, and a cut-off of 12 [ was applied for PME and also for van der Waals interactions. The default parameters of the g_mmpbsa tool were utilized for the calculations of the MM/PBSA binding energies [30]. To ensure stable binding conformations of the protein molecules or the ligands, no restraint or external forces were implemented on the system in case of the conventional MD simulation. The simulations were primarily done to study the dynamic character of the protein-ligand interaction, for a highly detailed atomistic analysis of the binding stability of the complexes. Each of the complexes were simulated in sets of three independent MD simulations.

### Cell Culture

TRAIL-sensitive breast cancer cell line MDA-MB-231 and TRAIL-resistant breast cancer cell line MCF7 and MDA-MB-453 were cultured in a high glucose DMEM medium supplemented with 10% fetal bovine serum (FBS) and 1% penicillin-streptomycin. The cells were maintained at 37°C in an environment with saturated humidity and 5% CO2. This study was approved by the Institutional Biosafety Committee of Presidency University in Kolkata, India. All cell culture reagents and plasticware were sourced from ThermoFisher Scientific, USA.

### MTT Assay

Live cells contain active NADPH-dependent oxidoreductase enzymes, which reduce the MTT dye, 3-(4,5-dimethylthiazol-2-yl)-2,5-diphenyltetrazolium bromide, into insoluble purple formazan crystals. In contrast, dead cells lack this enzymatic activity. Therefore, a deeper purple color signifies higher cell viability. In this study, cell viability was assessed using the MTT assay according to the protocol described by Ghosal et. al. [31]. Cells were seeded in a 96-well plate at a density of 50,000 cells per well. After the respective treatments, the culture media in each group was replaced with fresh media containing MTT (SRL, India) and incubated for 4 hours. The formazan crystals formed were dissolved in DMSO, and absorbance was measured at 590 nm using a SynergyH1 microplate reader with Gen5 software (Biotek, USA). All experiments were conducted in triplicate and repeated three times to ensure statistical accuracy.

### RNA Isolation

After cell harvesting, RNA was isolated using TRIzol reagent (Ambion, Life Technologies, USA). After adding TRIzol to the cell pellet, the mixture was vortexed at medium speed for 15 seconds and then incubated on ice for 10 minutes. The sample was centrifuged at 12,000 rpm for 10 minutes at 4°C, and the resultant aqueous phase was collected. Isopropanol was added, followed by another round of centrifugation at 12,000 rpm for 10 minutes at 4°C. The resulting visible RNA pellets were washed with 70% ethanol and then dissolved in DEPC-treated autoclaved water at 60°C. The quantity and quality of the RNA were evaluated using the A260/A280 absorbance ratio and agarose gel electrophoresis, respectively.

### Quantitative real-time PCR (qPCR) analysis

For transcript analysis of DR5 and DCR2, qPCR were performed using SsoFast Evagreen Supermix (Bio-Rad) on a PCR system 9700 (Bio-Rad), with β-actin as the endogenous control. The primer sequences used were as follows: for DR5, 5’-GGATCCACGTGAAGTCCTGT-3’ (forward) and 5’-TAACCCATCTGCCTCTGTCC-3’ (reverse); for β-actin, 5’-CCCAGCACAATGAAGATCAA-3’ (forward) and 5’-ACATCTGCTGGAAGGTGGAC-3’ (reverse); and for DCR2, 5’-GGTCATGGTGCAGGAACTTT-3’ (forward) and 5’-AGTGGAACTGGCAGCTGATT-3’ (reverse). The threshold cycle (CT) values were determined using CFX Manager software (Version 3.1.1517.0823) from Bio-Rad. Relative gene expression levels were calculated using the comparative CT method with β-actin as the reference gene. All experiments were conducted in duplicate and repeated three times.

### Trypan Blue Assay

The trypan blue dye exclusion test was employed to identify nonviable cells in the cytotoxicity assay. Here we employed the test to ascertain the capacity of different doses of paclitaxel to produce cytotoxicity in MDA-MB-231 and MCF7 cell lines. In summary, 5 × 10^4^ cells were incubated for 24-hours in a 24-well culture plate. Subsequent to different doses of paclitaxel administration for different timepoints, these cells were subjected to trypsinization and subsequently washed with PBS. Next, they were stained with trypan blue dye (Sigma-Aldrich, now part of Merck, Germany). The single-cell suspension was counted using a hemocytometer according to the method described by Berry et al. [32].

### Immunoblot Assay

Immunoblot analysis was used to check the expression of DR5 and DCR2 proteins after specific treatments. Cells were harvested using trypsin, and total protein was isolated using ready-to-use RIPA buffer according to the manufacturer’s instructions (Sigma-Aldrich, USA). Protein lysates were separated by their molecular weight using sodium dodecyl sulfate-polyacrylamide gel electrophoresis (SDS-PAGE) and then transferred onto a PVDF membrane (Millipore, Germany). Target proteins were identified using primary antibodies against DR5 (Cat. No.8074) and DCR2 (Cat. No.8049) from Cell Signaling Technology, USA. The endogenous control used was β-actin (Cat. No. BB-AB0024S, Bio Bharati Life Science, India). Protein bands were detected using an enhanced chemiluminescence (ECL) reagent (Bio-Rad, USA) and visualized with a gel documentation system (Bio-Rad, USA). Protein band intensities were quantified using Image J software and normalized to the endogenous control. Each experiment was conducted three times to ensure statistical validity.

### Colony Formation Assay

To assess clonogenic survival, MDA-MB-453 cells were harvested, counted, and seeded into 6-well plates (10^4^ cells per well) according to the experimental groups. Following attachment, cells were subjected to treatment and incubated at 37 °C in a humidified CO_2_ incubator for 3 weeks until colonies formed. After the appearance of visible colonies in the control well, the medium was removed, and the cells were rinsed twice with PBS. Thereafter the cells were fixed with 10% (v/v) neutral buffered formalin solution for 20 minutes. The formalin solution was thereafter removed, and the cells were stained with a 0.5% (v/v) crystal violet solution for 45 minutes. Colonies were subsequently enumerated utilizing a microscope and the colony counter plugin in Fiji software.

## Results

### In silico analysis shows paclitaxel to facilitate the binding of TRAIL with DR5 in comparison to its binding with DCR2

AutoDock vina results point to the highly stable nature of paclitaxel molecule in complex with the DCR2 or DR5 binding site along with TRAIL as is indicated by the high negative value of binding energy obtained from docking study (–7 kcal/mol for DCR2 and –6.3 kcal/mol for DR5 based on the best conformations). The best binding structures can be seen in supplementary fig 1. MD simulation analysis showed the highly stable nature of TRAIL-DR5 (fig. 1, left – upper panel) and TRAIL-DCR2 (fig.1, left – lower panel) complexes as is indicated by the low RMSD (root mean square value) values of the protein backbone atoms in both cases (0.33 ± 0.06 nm for TRAIL-DR5 complex and 0.48 ± 0.08 nm for TRAIL-DCR2 complex). Binding energy analysis shows that in absence of paclitaxel, TRAIL has more binding propensity towards DCR2 (binding energy for TRAIL-DCR2 complex: –688 KJ/mol) (fig.1, left – lower panel) in comparison to DR5 (binding energy for TRAIL-DR5 complex: –580 KJ/mol) (fig. 1, left – upper panel). Thus, when both the receptors are present, taking all other conditions to be constant it can be said that in absence of paclitaxel, TRAIL has a slightly more binding propensity towards DCR2 than DR5.

**Fig 1:**
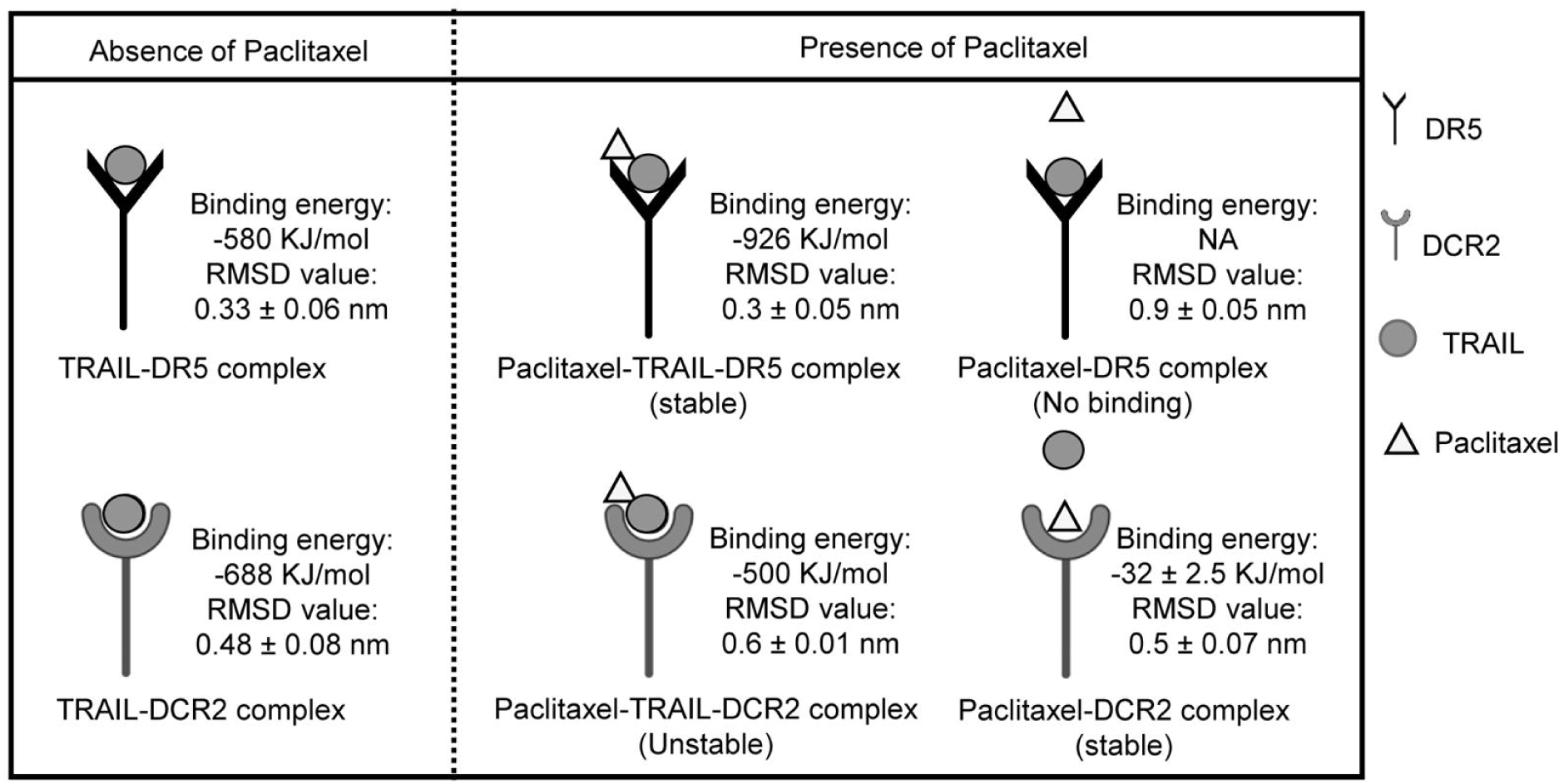
*In silico* findings suggest that paclitaxel plays a role in favoring the association of TRAIL with DR5 over DCR2. MM/PBSA binding energies and RMSD values for various molecular complexes: Left panels: TRAIL-DR5 complex (upper) and TRAIL-DCR2 complex (lower) in the absence of Paclitaxel. Middle panels: TRAIL-DR5 complex (upper) and TRAIL-DCR2 complex (lower) in the presence of paclitaxel. Right panels: Paclitaxel-DR5 complex (upper) and Paclitaxel-DCR2 complex (lower).

**Fig 2:**
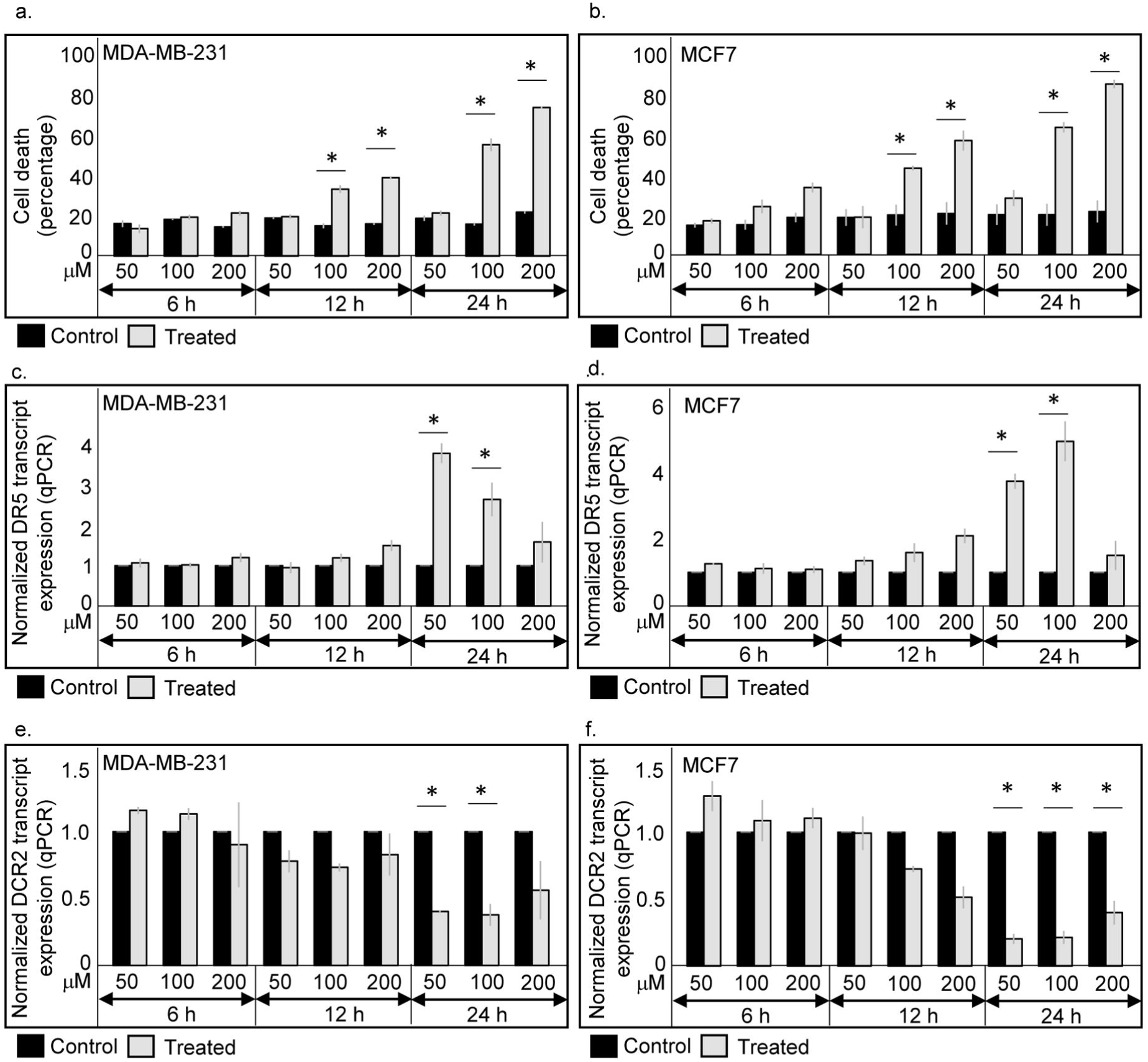
Paclitaxel administration results in elevated transcript level of DR5 and reduced transcript level of DCR2 in MDA-MB-231 and MCF7 cells. **a, b** Trypan blue assay showing percentage of cell death in MDA-MB-231 and MCF7 cells respectively after paclitaxel treatment at a dose of 50 nM, 100 nM and 200 nM for 6-hours, 12-hours and 24-hours. **c, d** Quantitative PCR showing transcript expression of DR5 in MDA-MB-231 and MCF7 cells respectively after paclitaxel treatment at a dose of 50 nM, 100 nM and 200 nM for 6-hours, 12-hours and 24-hours. **e, f** Quantitative PCR showing transcript expression of DCR2 in MDA-MB-231 and MCF7 cells respectively after paclitaxel treatment at a dose of 50 nM, 100 nM and 200 nM for 6-hours, 12-hours and 24-hours. Transcript analysis was done for DR5 and DCR2 in both cell lines through real time quantitative PCR (qPCR) using β-actin as an endogenous control. Graph represents mean expression value obtained from three biological repeats for every experiment. * Indicates p value ≤ 0.05.

Next, in presence of paclitaxel, RMSD value of the paclitaxel-TRAIL-DR5 (fig. 1, middle – upper panel) complex was seen to be 0.3 ± 0.05 nm indicating slightly more stabilization in comparison to TRAIL-DR5 complex (RMSD value: 0.33 ± 0.06 nm) (fig. 1, left – upper panel). The binding energy for paclitaxel-TRAIL-DR5 complex is –926 KJ/mol whereas the binding energy for TRAIL-DR5 complex was –580 KJ/mol thereby indicating a stabilizing effect of paclitaxel on TRAIL-DR5 interaction. In case of paclitaxel-DR5 (fig. 1, right – upper panel) complex, it was seen that the paclitaxel molecule got dissociated from DR5 indicating the inherent less stable binding between the DR5 molecule and the paclitaxel molecule. The RMSD value for paclitaxel-DR5 complex was seen to be 0.9 ± 0.5 nm indicating gross destabilization of the paclitaxel molecule in the DR5 protein binding pocket. In this case, no binding energy was found indicating that paclitaxel molecule got unbound from DR5 molecule. However, in presence of paclitaxel, the paclitaxel-TRAIL-DCR2 (fig. 1, middle – lower panel) complex was found to be stable in nature as indicated by the RMSD value i.e. 0.6 ± 0.1 nm. This higher RMSD value found in paclitaxel-TRAIL-DCR2 (fig. 1, middle – lower panel) complex in comparison to TRAIL-DCR2 complex (in absence of paclitaxel) (fig. 1, left – lower panel) indicates a destabilizing influence of paclitaxel molecule on TRAIL-DCR2 complex. The binding energy for paclitaxel-TRAIL-DCR2 complex is –500 KJ/mol (fig. 1, middle – lower panel) whereas the binding energy for TRAIL-DCR2 complex was –688 KJ/mol (fig. 1, left – lower panel) thereby indicating a destabilizing effect of paclitaxel on TRAIL-DCR2 interaction. The RMSD value of paclitaxel-DCR2 complex (fig. 1, right – lower panel) was found to be 0.5 ± 0.07 nm indicating the binding stability of paclitaxel with DCR2. The same can be ascertained from the binding energy of paclitaxel with the DCR2 binding pocket which came out to be – 32 ± 2.5 kJ/mol. Thus, from *in silico* analysis we can conclude that paclitaxel may selectively block DCR2, thereby facilitating stable interaction of DR5 with TRAIL.

### Paclitaxel treatment increases DR5 expression and decreases DCR2 expression in MDA-MB-231 and MCF7 cells at transcript levels

To investigate the effect of paclitaxel treatment on breast cancer cells, different concentrations of paclitaxel and exposure time points were first determined in MDA-MB-231 and MCF7 cell line. In our cell culture study, LD50 of paclitaxel for MDA-MB-231 and MCF7 was found to be 113.5 nM and 101.5 nM, respectively, at 24-hours. Therefore, three different dosages of 50 nM, 100 nM, and 200 nM of paclitaxel were chosen to look into the effects of varying drug concentrations on TRAIL-sensitive MDA-MB-231 and TRAIL-resistant MCF7 breast cancer cells. Trypan blue assay showed that there was significant cell death at 100 nM and 200 nM of paclitaxel administration for 12– and 24-hours’ time point both in MDA-MB-231 (fig.2a) and MCF7 cells (fig.2b). Next, we studied the transcript expression of DR5 and DCR2 at the above-mentioned doses. In MDA-MB-231 as well as MCF7 cells, DR5 transcript expression was found to be significantly increased after treating the cells with 50 nM and 100 nM dose of paclitaxel for 24-hours (fig.2c, 2d). Moreover, DCR2 transcript expression was decreased at the same dose of 50 nM and 100 nM after 24-hours paclitaxel treatment in MDA-MB-231 cells (fig.2e). In MCF7 cells, transcript expression of DCR2 was decreased significantly at 50 nM, 100 nM and 200 nM doses for 24-hours (fig.2f). Taken together, we can conclude that at 50 nM of paclitaxel treatment for 24-hours there was no significant cell death, but at the same time point DR5 transcript expression was increased whereas DCR2 transcript expression was decreased for both the cell lines.

### Paclitaxel treatment increases DR5 expression and decreases DCR2 expression in MDA-MB-231 and MCF7 cells at the protein level

Cell viability using MTT assay was performed to reconfirm the cell death percentage as obtained by trypan blue assay at 50 nM, 100 nM and 200 nM doses of paclitaxel for 24-hours. Both in MDA-MB-231 and MCF7 cells, significant decrease in cell viability was observed at 100 nM and 200 nM of paclitaxel administration for 24-hours’ time point (supplementary fig 2a, 2b). However, 50 nM paclitaxel administration did not decrease cell viability significantly. To eliminate high-concentration-induced drug-related toxicity, 100 nM and 200 nM dosages were eliminated from downstream experiments. A non-lethal paclitaxel dose of 50 nM at 24-hours of treatment was selected for further analysis for both the cell lines. Next, we analyzed the expression of DR5 and DCR2 proteins specifically at 50 nM dose of paclitaxel for 24-hours. TRAIL sensitive MDA-MB-231 cells showed significant increase in DR5 protein expression (fig. 3a) as well as significant decrease in DCR2 protein expression (fig. 3b) after treatment with non-toxic dose of paclitaxel. Protein expression of DR5 and DCR2 was also observed to be increased and decreased, respectively, in TRAIL-resistant MCF7 cell line (p<0.05) (fig. 3c, 3d) on paclitaxel treatment. Thus, we can conclude that treatment with a non-toxic dose of paclitaxel led to an increase in DR5 protein expression, along with a decrease in DCR2 protein expression in both MDA-MB-231 and MCF7 cells.

**Fig 3:**
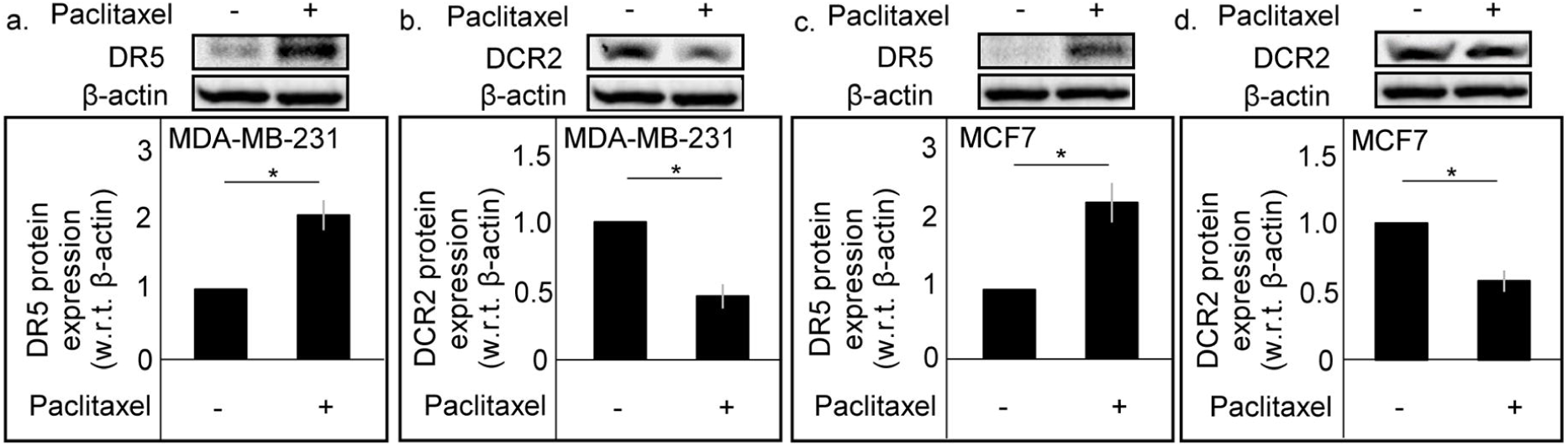
Treatment with Paclitaxel augmented DR5 protein expression while diminishing DCR2 protein expression in MDA-MB-231 and MCF7 cell lines. **a, b** Immunoblots showing protein expression of DR5 and DCR2 respectively in MDA-MB-231 cells after paclitaxel treatment at a dose of 50 nM for 24 hours. **c, d** Immunoblots showing protein expression of DR5 and DCR2 respectively in MCF7 cells after paclitaxel treatment at a dose of 50 nM for 24 hours. β-actin is used as an endogenous control. Densitometric graph represents mean expression value obtained from three biological repeats of every experiment whereas blot shows the representative image. * Indicates p value ≤ 0.05.

### Pre-treatment with Paclitaxel elicits rhTRAIL mediated cell death in TRAIL-resistant cell lines MCF7 and MDA-MB-453

Since administering paclitaxel at a dose of 50 nM for 24-hours elevated DR5 expression and decreased DCR2 expression but did not cause any significant cell death in TRAIL-sensitive MDA-MB-231 and TRAIL-resistant MCF7 cells, we investigated the functional significance of sequential treatment of paclitaxel followed by TRAIL treatment in TRAIL-resistant MCF7 and MDA-MB-453 cells. Four experimental groups were designed, group 1 – control, group 2 – treated with 50 nM of paclitaxel for 24-hours only, group 3 – treated with 250 ng/ml of rhTRAIL for 6-hours only and group 4 – treated with 50 nM paclitaxel for 24-hours and then after media change, 250 ng/ml rhTRAIL was administered for a duration of 6-hours (supplementary fig 3). MTT assay was carried out for the above-mentioned experimental groups to determine the sensitivity of TRAIL-resistant cell lines to sequential drug treatment. Exposure to 50 nM paclitaxel for 24-hours followed by TRAIL treatment for 6-hours caused significant decrease in cell viability in group 4 in comparison to the other three experimental groups both in MCF7 (fig. 4a) and MDA-MB-453 (fig. 4b) cells. Furthermore, immunoblot data showed significantly increased expression of cleaved caspase 3 in group 4 compared to other groups, both in MCF7 (fig. 4c) and MDA-MB-453 (fig. 4d) cells. Therefore, we can conclude that pre-treatment of paclitaxel elicits rhTRAIL mediated cell death in TRAIL-resistant cells.

**Fig 4:**
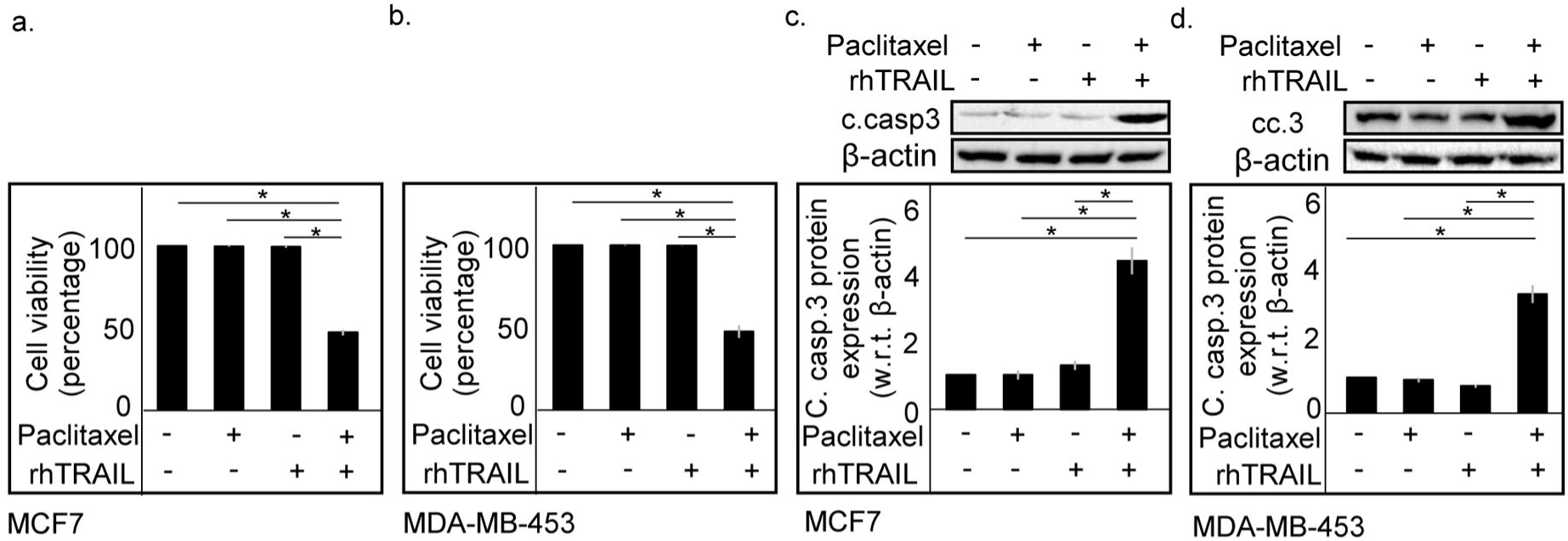
Pre-treatment with paclitaxel sensitized the TRAIL-resistant human breast cancer cell lines MCF7 and MDA-MB-453 to rhTRAIL mediated cell death. **a, b** bar diagram showing percentage of cell viability (MTT assay) in MCF7 and MDA-MB-453 cells respectively after pre-treatment with paclitaxel followed by rhTRAIL treatment. Bar graphs represent mean value obtained from three biological repeats of each respective experiments. **c, d** immunoblots showing protein expression of cleaved caspase-3 in MCF7 and MDA-MB-453 cells respectively after the treatment protocol. β-actin is used as an endogenous control. Densitometric graph represents mean expression value obtained from three biological repeats of every experiment whereas blot shows the representative image. * Indicates p value ≤ 0.05.

### Pre-treatment with paclitaxel decreases the survival of TRAIL-resistant MDA-MB-453 cells upon rhTRAIL treatment

A colony formation assay was conducted to investigate the long-term effects of the combination treatment on TRAIL resistant cells. Exposure to 50 nM paclitaxel for 24 hours, followed by rhTRAIL treatment for 6 hours, resulted in a marked reduction in cell survival in group 4 compared to the other three experimental groups in MDA-MB-453 cells (fig. 5a, b). No significant decrease in colonies were noted in either the paclitaxel-treated group or the rhTRAIL-treated group relative to the control group. This finding demonstrates that pretreatment with a sub-lethal dose of paclitaxel decreases the survival of TRAIL-resistant cells upon rhTRAIL treatment.

**Fig 5:**
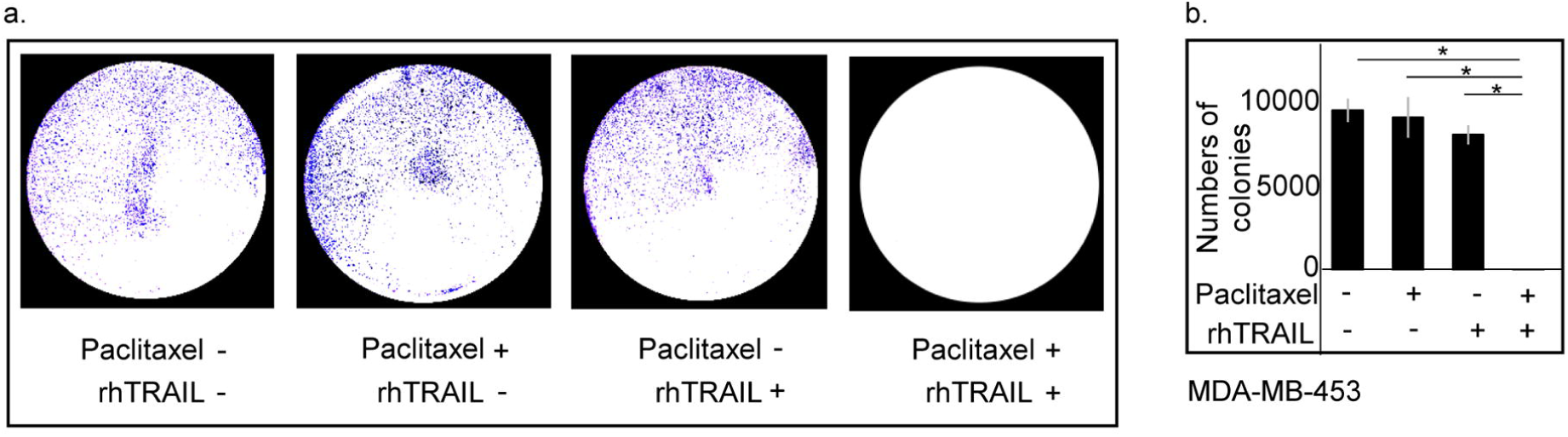
Pre-treatment with paclitaxel decreases the survival of TRAIL-resistant MDA-MB-453 cells upon rhTRAIL treatment. **a** photograph showing number of colonies in MDA-MB-453 cells after pre-treatment with paclitaxel followed by rhTRAIL treatment. b. Bar graphs represent the average number of colonies from three biological repeats. * Indicates p value ≤ 0.05.

### Pre-treatment with Paclitaxel sensitizes TRAIL-resistant cell lines MCF7 and MDA-MB-453 to rhTRAIL-induced cell death by upregulation of DR5 and downregulation of DCR2

To explore the reason behind how paclitaxel pre-treatment increases rhTRAIL mediated apoptosis in TRAIL-resistant cells, effect of paclitaxel treatment on DR5 and DCR2 protein expression was checked. Immunoblot analysis revealed a significant increase in DR5 protein expression and significant decrease in DCR2 protein expression in paclitaxel alone group as well as in that treated with both paclitaxel and TRAIL in comparison to the control group, both in MCF7 and MDA-MB-453 (fig. 6a, 6b) cells. So, we can conclude that pre-treatment with paclitaxel increases the expression of DR5 and decrease the expression of DCR2 in MCF7 and MDA-MB-453 thereby making them sensitive to rhTRAIL mediated apoptosis.

**Fig 6:**
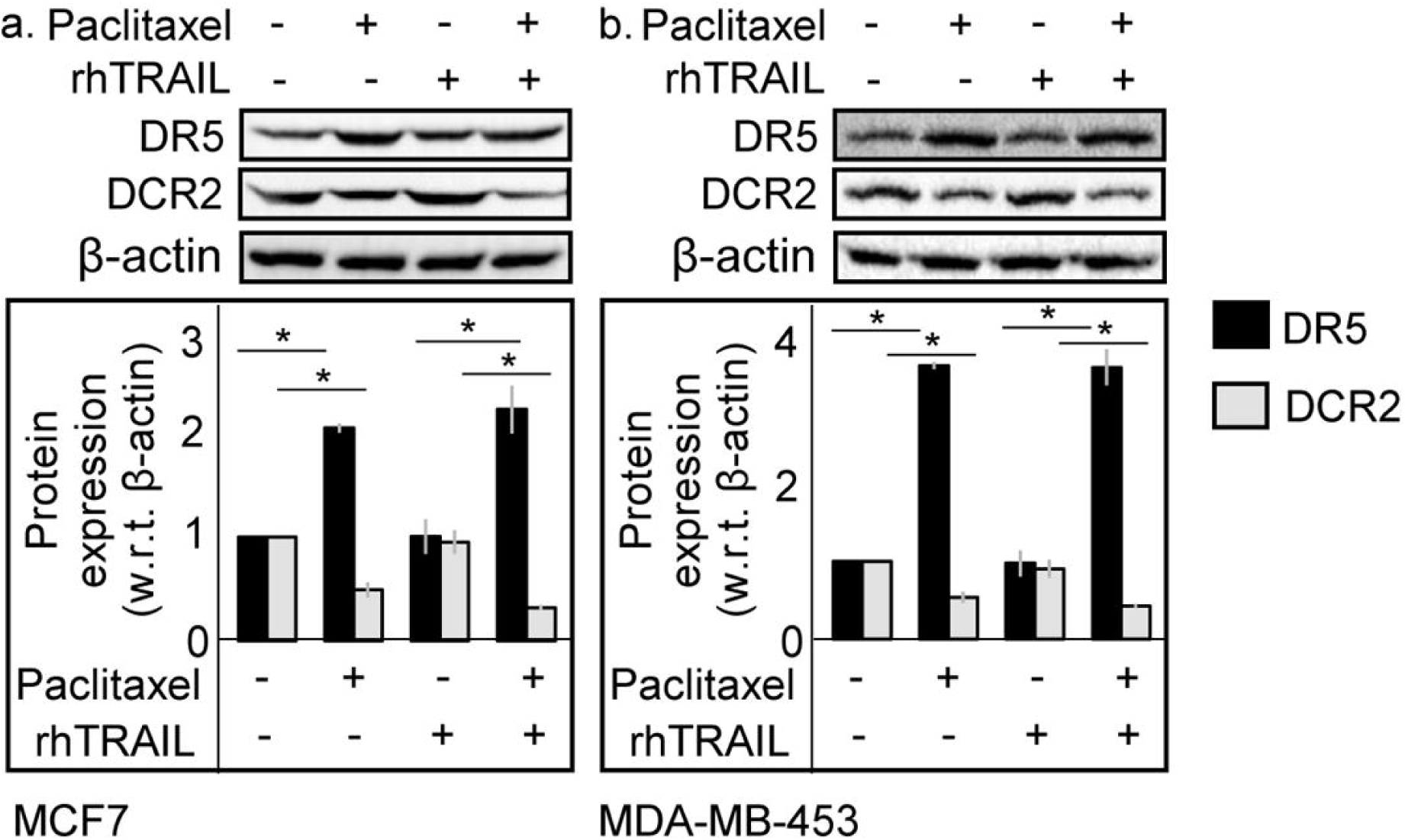
Paclitaxel pre-treatment sensitizes the TRAIL-resistant human breast cancer cell lines MCF7 and MDA-MB-453 to rhTRAIL induced cell death via the upregulation of DR5 and downregulation of DCR2. **a** Immunoblot showing protein expression of DR5 and DCR2 in MCF7 cells after the treatment protocol. **b** Immunoblots showing protein expression of DR5 and DCR2 in MDA-MB-453 cells after the treatment protocol, β-actin is used as an endogenous control. Densitometric graph represents mean expression value obtained from three biological repeats of every experiment whereas blot shows the representative image. * Indicates p value ≤ 0.05.

## Discussion

Tumor Necrosis Factor (TNF) superfamily can trigger cell death through interactions between TNF ligands and their cognate death receptors. However, majority of the pro-apoptotic ligands, kill healthy cells. rather than targeting tumors specifically [33]. Since studies have demonstrated that TRAIL, a member of the TNF superfamily, can kill cancerous cells while sparing healthy cells, it has gained attention as a promising potential anti-cancer protein [34, 35]. However, some cancer cells are inherently resistant to TRAIL-mediated apoptosis [36]. Aberrant signaling via death or decoy receptors is one of the mechanisms by which TRAIL resistance can occur. By binding and sequestering TRAIL, decoy receptors prevent the signaling cascade that causes death. Resistance to TRAIL can therefore be conferred by either increased expression of the decoy receptors or, on the other hand, decreased expression of the death receptors [3]. From our earlier study we have observed that TRAIL-resistant cells show increased expression of DCR2 relative to DR5 [31]. Therefore, therapeutic approaches that aim to block or downregulate decoy receptors serve as a way to counteract TRAIL resistance while simultaneously enhancing the capacity of the drug to induce apoptosis by increasing TRAIL-death receptor interactions. The potent anti-cancer agent paclitaxel binds to the beta-tubulin subunit of microtubules and thus leads to inhibition of mitotic spindle formation, ultimately resulting in cellular apoptosis [37]. However, it has been shown that paclitaxel can also impacting cancer cell proliferation and growth by inhibiting tumor angiogenesis and inducing the expression of genes and cytokines that suppress cell growth and trigger apoptosis [37, 38]. Several studies have shown paclitaxel and TRAIL combinatorial treatment to result in a significantly enhanced tumoricidal potential in non-small cell lung cancer, ovarian cancer, gastric cancer, melanoma and prostate cancer [23, 39, 40]. Furthermore, paclitaxel-induced apoptosis has been associated with increased levels of death receptor proteins in prostate gland cells, renal cancer cells, and gliomas [23, 41]. Therefore, our aim was to investigate the role of paclitaxel in sensitizing the TRAIL-resistant breast cancer cells towards TRAIL-mediated apoptosis.

At first, *in silico* modelling and simulation was done to investigate the *in-silico* interaction between TRAIL, DR5, DCR2 and paclitaxel with each other. Docking study indicated highly stable nature of paclitaxel molecule in complex with the DCR2 or DR5 binding site along with TRAIL (supplementary fig 1). From the MD simulation analysis, it was found out that, in absence of paclitaxel, binding energy for TRAIL-DR5 complex was –580 KJ/mol and RMSD value was 0.33 ± 0.06 nm. On the other hand, binding energy for TRAIL-DCR2 complex was –688 KJ/mol and RMSD value was 0.48 ± 0.08 nm. This shows that in absence of paclitaxel, TRAIL has more binding affinity towards DCR2 in comparison to DR5. Next, we have checked the interaction between paclitaxel alone with DR5 or DCR2 receptor. In case of the paclitaxel-DR5 complex, it was seen that the paclitaxel molecule does not bind with DR5 as the RMSD value (0.9 ± 0.5 nm) for paclitaxel-DR5 complex is quite high. This indicates gross destabilization of the paclitaxel molecule from the DR5 protein binding pocket. Added to this, no binding energy was found between DR5 and paclitaxel molecule indicating that paclitaxel molecule remains dissociated from DR5 molecule. In case of the paclitaxel-DCR2 complex, it was seen that the paclitaxel molecule bind spontaneously with DCR2 as the RMSD value of paclitaxel-DCR2 complex was found to be 0.5 ± 0.07 nm and binding energy of paclitaxel with DCR2 was –32 ± 2.5 kJ/mol, indicating the binding stability. This reflects that paclitaxel molecule can specifically block DCR2 ligand binding site without interrupting DR5. Next, the interactions between paclitaxel and TRAIL-DR5 complex or TRAIL-DCR2 complex were explored. In presence of paclitaxel, the RMSD value of TRAIL-DCR2 complex (fig 1d) was 0.6 ± 0.1 nm and the binding energy was –500 KJ/mol. This higher RMSD value and a decrease in negative binding energy between TRAIL and DCR2 in presence of paclitaxel compared to these values obtained in absence of paclitaxel, indicates a destabilizing influence of paclitaxel molecule on TRAIL-DCR2 complex. Next, in presence of paclitaxel, RMSD value of the TRAIL-DR5 complex was seen to be 0.3 ± 0.05 nm and the binding energy was –926 KJ/mol. This slightly lower RMSD value and a large increase in negative binding energy between TRAIL and DR5 in presence of paclitaxel compared to these values obtained in absence of paclitaxel, indicates a stabilizing influence of paclitaxel molecule on TRAIL-DR5 complex. Thus, it can be seen that in absence of paclitaxel molecule a preference towards DCR2 was seen for TRAIL molecule in comparison to DR5 protein. Next, the binding of paclitaxel was found to be stable in case of DCR2 (fig 1f) while it was unstable in case of DR5 indicating that paclitaxel would preferentially bind with high stability to DCR2 and not DR5. Lastly, presence of paclitaxel molecule led to a destabilization of the TRAIL-DCR2 complex while it further stabilized the TRAIL-DR5 molecule changing the preference of the TRAIL molecule towards DR5 in comparison to DCR2. From our *in-silico* study we conclude that paclitaxel exclusively binds and blocks the DCR2 receptor (fig 1).

Taking notions from *in silico* study that paclitaxel can be an effective blocker of DCR2, we shifted to *in vitro* study to explore the mechanism of paclitaxel mediated cell death in absence or presence of TRAIL and their overall implication in breast cancer cells. To study the cellular effects of varying paclitaxel concentrations, doses of 50 nM, 100 nM, and 200 nM were tested over 6-, 12-, and 24-hours. Our aim was to evaluate the effect of paclitaxel on cell death in TRAIL sensitive MDA-MB-231 and TRAIL-resistant MCF7 cell line. Significant cell death was observed at 100 nM and 200 nM of paclitaxel administration for 12– and 24-hours’ time point in both the MDA-MB-231 and MCF7 cell lines. However, 50 nM dose of paclitaxel did not elicit cell death at any of the time points in both the breast cancer cells (fig 2a, 2b). This data was validated using MTT assay where it was confirmed that 50 nM dose of paclitaxel at 24-hours’ time point did not alter cell viability in all the three breast cancer cell lines (supplementary fig 2a, 2b, 2c). However, we observed DR5 transcript expression to increase in MDA-MB-231 and MCF7 cell lines at this non-toxic dose of paclitaxel (fig 2c, 2d). Moreover, DCR2 transcript expression was also found to be decreased in these breast cancer cells (fig 2e, 2f). Furthermore, non-lethal dose of paclitaxel administration caused increased DR5 and decreased DCR2 protein expression in both the TRAIL sensitive and TRAIL-resistant breast cancer cell lines (fig 3a, 3b, 3c, 3d).

As administration of paclitaxel can increase DR5 expression and decrease DCR2 expression, we hypothesized that pretreatment of paclitaxel can sensitize the TRAIL-resistant cells towards TRAIL-mediated apoptosis. Therefore, we used two TRAIL-resistant cell lines, MCF-7 and MDA-MB-453 for further study. Cell viability assay showed that pretreatment of paclitaxel followed by rhTRAIL treatment led to significantly more cell death in TRAIL-resistant MCF7 and MDA-MB-453 cell lines compared to the paclitaxel alone or TRAIL alone treated groups (fig 4a, 4b). Additionally, the combinatorial treatment resulted in higher levels of cleaved caspase 3 in both the cell lines compared to the paclitaxel alone or TRAIL alone treated groups, indicating that pre-treatment with paclitaxel sensitizes these TRAIL-resistant breast cancer cells towards rhTRAIL-mediated apoptosis. Moreover, colony formation assay showed that pre-treatment with paclitaxel decreases the long term survival of TRAIL-resistant MDA-MB-453 cells upon rhTRAIL administration (fig 5a, 5b.). To identify the cause of enhanced rhTRAIL sensitivity we investigated the expression of TRAIL receptors on combination therapy. We observed that paclitaxel pretreatment along with administration of rhTRAIL significantly increased the expression of pro-apoptotic DR5 protein and decreased the expression of anti-apoptotic DCR2 protein compared to the control in TRAIL-resistant MCF7 and MDA-MB-453 cell lines, thereby sensitizing TRAIL-resistant cells towards rhTRAIL (fig 6a, 6b).

Although TRAIL therapy comes across as an effective anti-cancer regimen, the major obstacle in its pre-clinical applications has been the problem of TRAIL resistance [42]. Through our study, we show that in breast tumors that are TRAIL-resistant in nature, engaging TRAIL therapy in association with paclitaxel pre-treatment can potentially lead to significant enhancement of apoptosis through the modulation of TRAIL receptor dynamics (fig 7).

**Fig 7:**
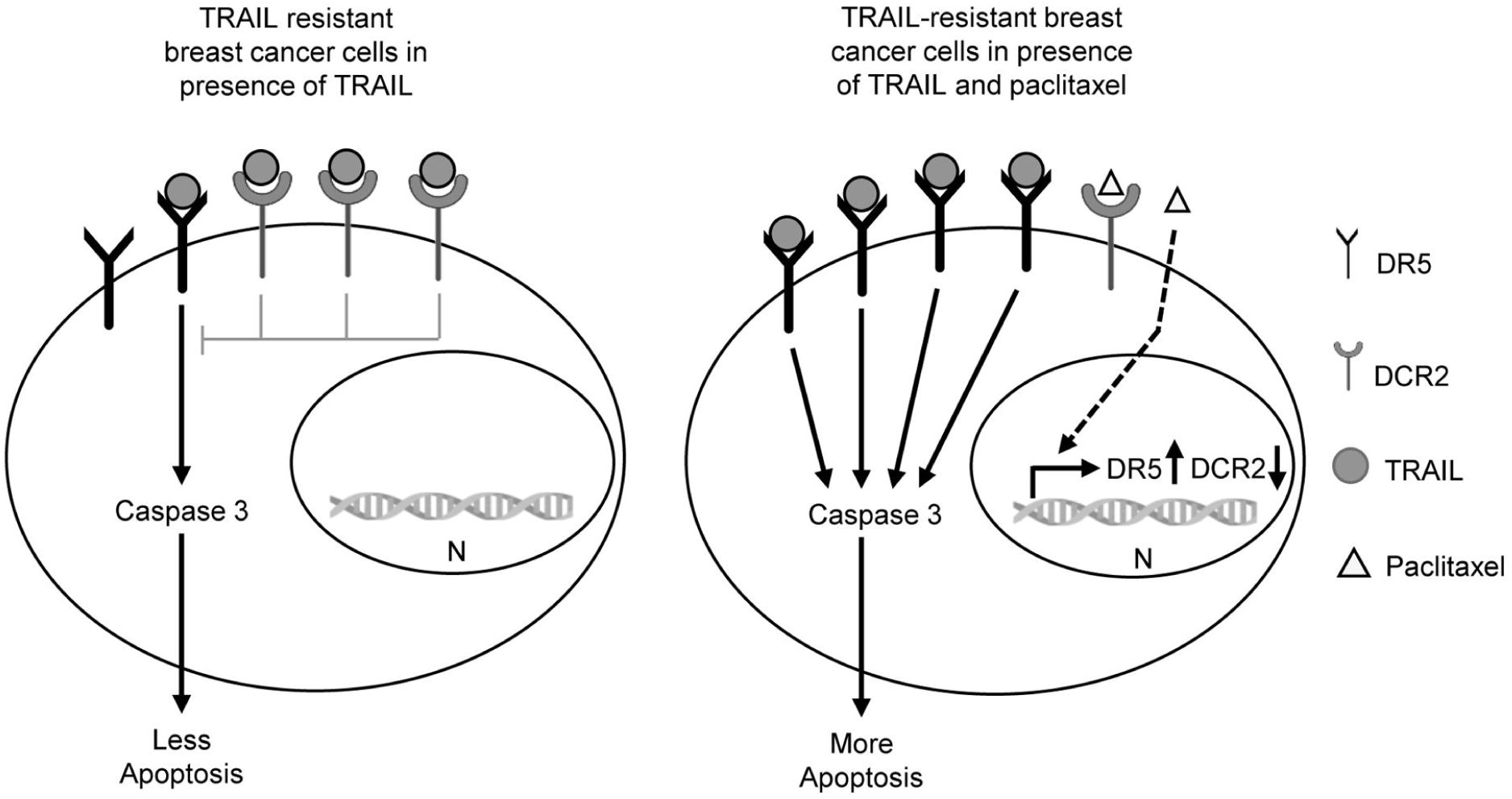
Schematic illustration depicting the proposed mechanism by which paclitaxel reverses TRAIL resistance in breast cancer cells. TRAIL on binding to DR5 activates apoptosis via caspase-3. However, TRAIL-DCR2 interaction inhibits TRAIL-mediated apoptosis. Typically, TRAIL-resistant cells exhibit higher expression levels of DCR2 relative to DR5. In the absence of paclitaxel, TRAIL exhibits a greater binding affinity for the DCR2 receptor. In these resistant cells, on administration of TRAIL, the ligand (TRAIL) binds to DCR2 and suppresses apoptosis. In the presence of paclitaxel, the expression of DR5 increases while the expression of DCR2 decreases through an unidentified mechanism. Additionally, paclitaxel can physically interact with the DCR2 receptor, hence enhancing TRAIL-DR5 binding and rendering TRAIL-resistant cells susceptible to TRAIL mediated apoptosis.

## Conclusion

Our findings suggest that pre-treatment with paclitaxel increases the expression of death receptor 5 and decreases the expression of decoy receptor 2 which sensitizes the TRAIL-resistant breast cancer cells towards rhTRAIL therapy. An important direction for future research would be the optimization of this concurrent treatment regimen in animal studies and clinical trials, thus aiming to develop novel therapeutic protocols that maximize the effectiveness of TRAIL based therapy.

## Acknowledgement

This work was supported by DBT. N.G. and P.T. were supported by fellowship from CSIR-NET, and DST-INSPIRE, respectively. The MDA-MB-453 breast cancer cell lines were a kind gift from Dr. Nabendu Biswas of Presidency University, Kolkata, India. Infrastructure support was made possible with funds from Presidency University, DBT-BUILDER and DST-FIST.

## Ethics approval

This study does not require ethics approval since no human or animal subjects were used for the experiment.

## Supplementary figure legend

**Supplementary fig 1:** Left: TRAIL (red) in complex with DR5 (blue) following 50 ns MD simulation bound to paclitaxel. Right: TRAIL (red) in complex with DCR2 (blue) following 50 ns MD simulation bound to paclitaxel. Paclitaxel has been shown using stick model.

**Supplementary fig 2:** **a, b, c** MTT assay showing percentage of cell viability in MDA-MB-231,MCF7 and MDA-MB-453 cells respectively after paclitaxel treatment at a dose of 50 nM, 100 nM and 200 nM for 24-hours. Bar graphs represent mean value obtained from three biological repeats of each respective experiment.

**Supplementary fig 3:** Schematic diagram of treatment protocol. Pre-treatment of paclitaxel was done at a dose of 50 nM for 24-hours. Next, after the media change, rhTRAIL treatment was done at a sub-lethal dose of 250 ng/ml for 6-hours.

## Reference

1. Siegel RL, Giaquinto AN, Jemal A. Cancer statistics, 2024. CA Cancer J Clin. 2024; 74: 12–49.

2. Johnstone RW, Frew AJ, Smyth MJ. The TRAIL apoptotic pathway in cancer onset, progression and therapy. Nat Rev Cancer. 2008; 8: 782–98.

3. Thorburn A, Behbakht K, Ford H. TRAIL receptor-targeted therapeutics: resistance mechanisms and strategies to avoid them. Drug Resist Updat. 2008; 11: 17–24.

4. Falschlehner C, Emmerich CH, Gerlach B, Walczak H. TRAIL signalling: decisions between life and death. Int J Biochem Cell Biol. 2007; 39: 1462–75.

5. Degli-Esposti MA, Dougall WC, Smolak PJ, Waugh JY, Smith CA, Goodwin RG. The novel receptor TRAIL-R4 induces NF-kappaB and protects against TRAIL-mediated apoptosis, yet retains an incomplete death domain. Immunity. 1997; 7: 813–20.

6. Ehrhardt H, Fulda S, Schmid I, Hiscott J, Debatin KM, Jeremias I. TRAIL induced survival and proliferation in cancer cells resistant towards TRAIL-induced apoptosis mediated by NF-kappaB. Oncogene. 2003; 22: 3842–52.

7. Lalaoui N, Morle A, Merino D, Jacquemin G, Iessi E, Morizot A, et al. TRAIL-R4 promotes tumor growth and resistance to apoptosis in cervical carcinoma HeLa cells through AKT. PLoS One. 2011; 6: e19679.

8. Morel J, Audo R, Hahne M, Combe B. Tumor necrosis factor-related apoptosis-inducing ligand (TRAIL) induces rheumatoid arthritis synovial fibroblast proliferation through mitogen-activated protein kinases and phosphatidylinositol 3-kinase/Akt. J Biol Chem. 2005; 280: 15709–18.

9. Nguyen PT, Nguyen D, Chea C, Miyauchi M, Fujii M, Takata T. Interaction between N-cadherin and decoy receptor-2 regulates apoptosis in head and neck cancer. Oncotarget. 2018; 9: 31516–30.

10. Vindrieux D, Reveiller M, Chantepie J, Yakoub S, Deschildre C, Ruffion A, et al. Down-regulation of DcR2 sensitizes androgen-dependent prostate cancer LNCaP cells to TRAIL-induced apoptosis. Cancer Cell Int. 2011; 11: 42.

11. Zhan C, Wei X, Qian J, Feng L, Zhu J, Lu W. Co-delivery of TRAIL gene enhances the anti-glioblastoma effect of paclitaxel in vitro and in vivo. J Control Release. 2012; 160: 630–6.

12. Dorsey JF, Mintz A, Tian X, Dowling ML, Plastaras JP, Dicker DT, et al. Tumor necrosis factor-related apoptosis-inducing ligand (TRAIL) and paclitaxel have cooperative in vivo effects against glioblastoma multiforme cells. Mol Cancer Ther. 2009; 8: 3285–95.

13. Di Cristofano F, George A, Tajiknia V, Ghandali M, Wu L, Zhang Y, et al. Therapeutic targeting of TRAIL death receptors. Biochem Soc Trans. 2023; 51: 57–70.

14. Ko JH, Sethi G, Um JY, Shanmugam MK, Arfuso F, Kumar AP, et al. The Role of Resveratrol in Cancer Therapy. Int J Mol Sci. 2017; 18.

15. Walsh V, Goodman J. From taxol to Taxol: the changing identities and ownership of an anti-cancer drug. Med Anthropol. 2002; 21: 307–36.

16. Sati P, Sharma E, Dhyani P, Attri DC, Rana R, Kiyekbayeva L, et al. Paclitaxel and its semi-synthetic derivatives: comprehensive insights into chemical structure, mechanisms of action, and anticancer properties. Eur J Med Res. 2024; 29: 90.

17. Rowinsky EK. The development and clinical utility of the taxane class of antimicrotubule chemotherapy agents. Annu Rev Med. 1997; 48: 353–74.

18. Wang TH, Wang HS, Soong YK. Paclitaxel-induced cell death: where the cell cycle and apoptosis come together. Cancer. 2000; 88: 2619–28.

19. Patt D, Gauthier M, Giordano S. Paclitaxel in breast cancer. Womens Health (Lond). 2006; 2: 11–21.

20. Moliterni A, Tarenzi E, Capri G, Terenziani M, Bertuzzi A, Grasselli G, et al. Pilot study of primary chemotherapy with doxorubicin plus paclitaxel in women with locally advanced or operable breast cancer. Semin Oncol. 1997; 24: S17-0-S-4.

21. Nimmanapalli R, Perkins CL, Orlando M, O’Bryan E, Nguyen D, Bhalla KN. Pretreatment with paclitaxel enhances apo-2 ligand/tumor necrosis factor-related apoptosis-inducing ligand-induced apoptosis of prostate cancer cells by inducing death receptors 4 and 5 protein levels. Cancer Res. 2001; 61: 759–63.

22. Vignati S, Codegoni A, Polato F, Broggini M. Trail activity in human ovarian cancer cells: potentiation of the action of cytotoxic drugs. Eur J Cancer. 2002; 38: 177–83.

23. Odoux C, Albers A. Additive effects of TRAIL and paclitaxel on cancer cells: implications for advances in cancer therapy. Vitam Horm. 2004; 67: 385–407.

24. Lin T, Zhang L, Davis J, Gu J, Nishizaki M, Ji L, et al. Combination of TRAIL gene therapy and chemotherapy enhances antitumor and antimetastasis effects in chemosensitive and chemoresistant breast cancers. Mol Ther. 2003; 8: 441–8.

25. Keane MM, Ettenberg SA, Nau MM, Russell EK, Lipkowitz S. Chemotherapy augments TRAIL-induced apoptosis in breast cell lines. Cancer Res. 1999; 59: 734–41.

26. Singh TR, Shankar S, Chen X, Asim M, Srivastava RK. Synergistic interactions of chemotherapeutic drugs and tumor necrosis factor-related apoptosis-inducing ligand/Apo-2 ligand on apoptosis and on regression of breast carcinoma in vivo. Cancer Res. 2003; 63: 5390–400.

27. Buchsbaum DJ, Zhou T, Grizzle WE, Oliver PG, Hammond CJ, Zhang S, et al. Antitumor efficacy of TRA-8 anti-DR5 monoclonal antibody alone or in combination with chemotherapy and/or radiation therapy in a human breast cancer model. Clin Cancer Res. 2003; 9: 3731–41.

28. Trott O, Olson AJ. AutoDock Vina: improving the speed and accuracy of docking with a new scoring function, efficient optimization, and multithreading. J Comput Chem. 2010; 31: 455–61.

29. Van Der Spoel D, Lindahl E, Hess B, Groenhof G, Mark AE, Berendsen HJ. GROMACS: fast, flexible, and free. J Comput Chem. 2005; 26: 1701–18.

30. Kumari R, Kumar R, Open Source Drug Discovery C, Lynn A. g_mmpbsa--a GROMACS tool for high-throughput MM-PBSA calculations. J Chem Inf Model. 2014; 54: 1951–62.

31. Ghosal N, Tapadar P, Biswas D, Pal R. ELF3 plays an important role in defining TRAIL sensitivity in breast cancer by modulating the expression of decoy receptor 2 (DCR2). Mol Biol Rep. 2024; 51: 671.

32. Berry JM, Huebner E, Butler M. The crystal violet nuclei staining technique leads to anomalous results in monitoring mammalian cell cultures. Cytotechnology. 1996; 21: 73–80.

33. Bremer E. Targeting of the tumor necrosis factor receptor superfamily for cancer immunotherapy. ISRN Oncol. 2013; 2013: 371854.

34. Ashkenazi A, Pai RC, Fong S, Leung S, Lawrence DA, Marsters SA, et al. Safety and antitumor activity of recombinant soluble Apo2 ligand. J Clin Invest. 1999; 104: 155–62.

35. Walczak H, Miller RE, Ariail K, Gliniak B, Griffith TS, Kubin M, et al. Tumoricidal activity of tumor necrosis factor-related apoptosis-inducing ligand in vivo. Nat Med. 1999; 5: 157–63.

36. Tapadar P, Pal A, Ghosal N, Kumar B, Paul T, Biswas N, et al. CDH1 overexpression sensitizes TRAIL resistant breast cancer cells towards rhTRAIL induced apoptosis. Mol Biol Rep. 2023; 50: 7283–94.

37. Abu Samaan TM, Samec M, Liskova A, Kubatka P, Busselberg D. Paclitaxel’s Mechanistic and Clinical Effects on Breast Cancer. Biomolecules. 2019; 9.

38. Taghian AG, Abi-Raad R, Assaad SI, Casty A, Ancukiewicz M, Yeh E, et al. Paclitaxel decreases the interstitial fluid pressure and improves oxygenation in breast cancers in patients treated with neoadjuvant chemotherapy: clinical implications. J Clin Oncol. 2005; 23: 1951–61.

39. Huang S, Zhang Y, Wang L, Liu W, Xiao L, Lin Q, et al. Improved melanoma suppression with target-delivered TRAIL and Paclitaxel by a multifunctional nanocarrier. J Control Release. 2020; 325: 10–24.

40. Li L, Wen XZ, Bu ZD, Cheng XJ, Xing XF, Wang XH, et al. Paclitaxel enhances tumoricidal potential of TRAIL via inhibition of MAPK in resistant gastric cancer cells. Oncol Rep. 2016; 35: 3009–17.

41. Gong J, Yang D, Kohanim S, Humphreys R, Broemeling L, Kurzrock R. Novel in vivo imaging shows up-regulation of death receptors by paclitaxel and correlates with enhanced antitumor effects of receptor agonist antibodies. Mol Cancer Ther. 2006; 5: 2991–3000.

42. Deng D, Shah K. TRAIL of Hope Meeting Resistance in Cancer. Trends Cancer. 2020; 6: 989–1001.

