## Supplementary figures and images for "Paclitaxel sensitizes TRAIL (tumor necrosis factor-related apoptosis-inducing ligand)-resistant breast cancer cells towards TRAIL-mediated apoptosis"

### Supplementary figure 1

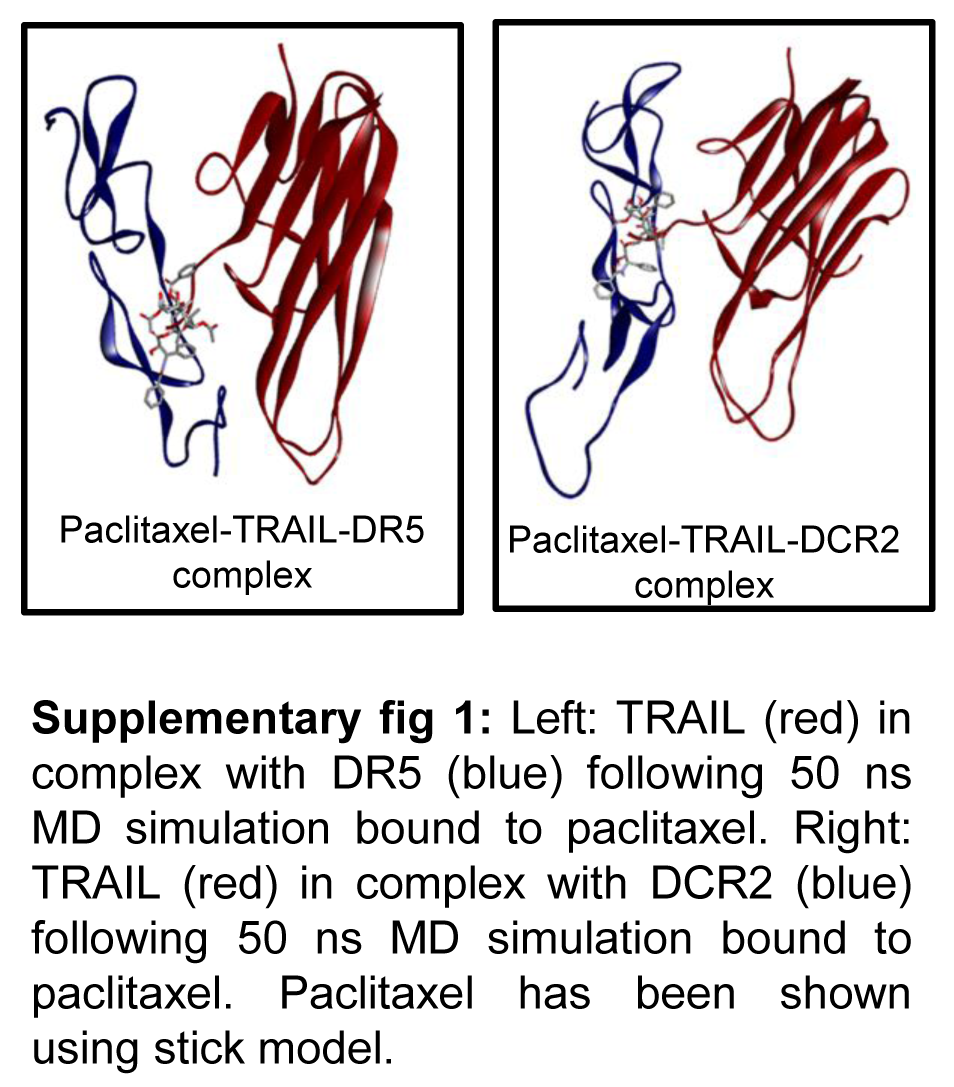

### Supplementary figure 2

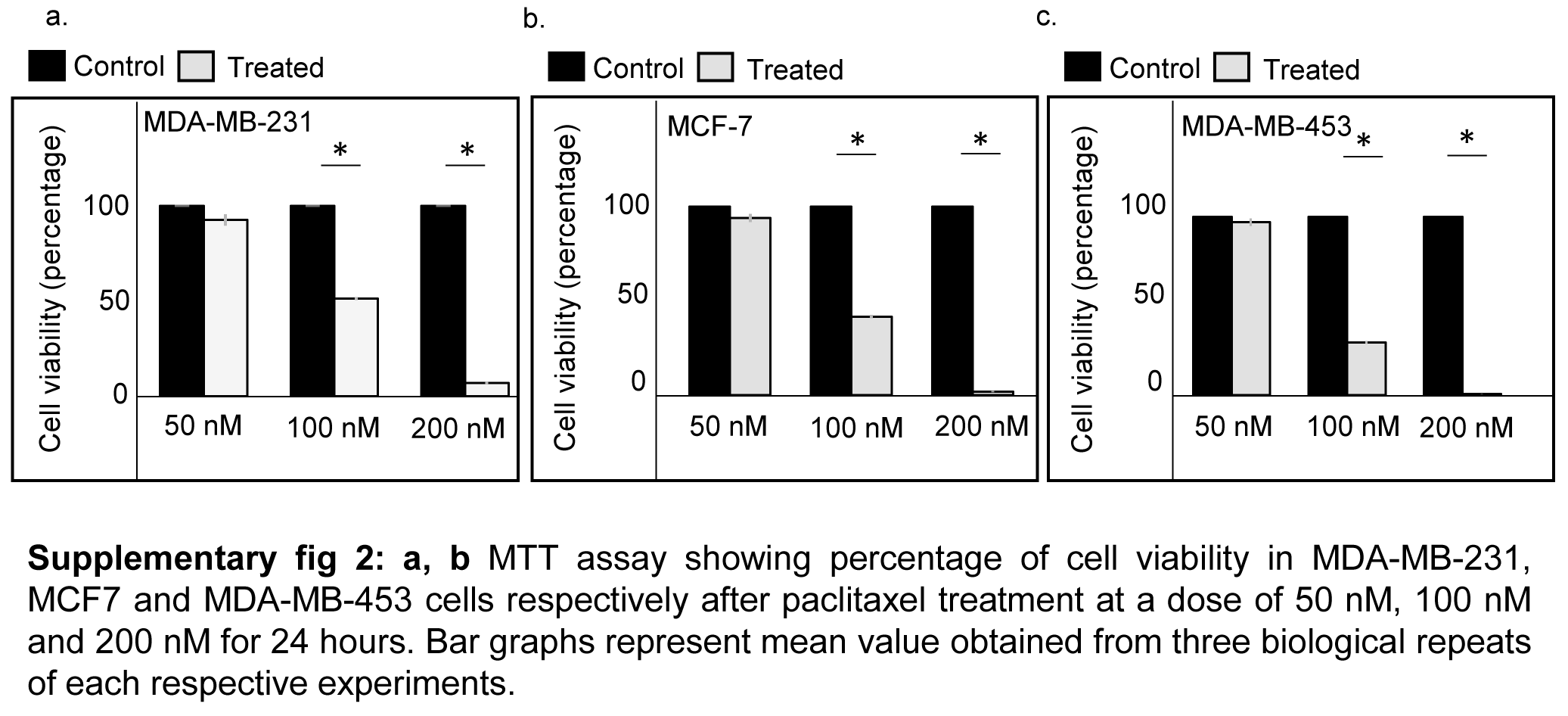

### Supplementary figure 3

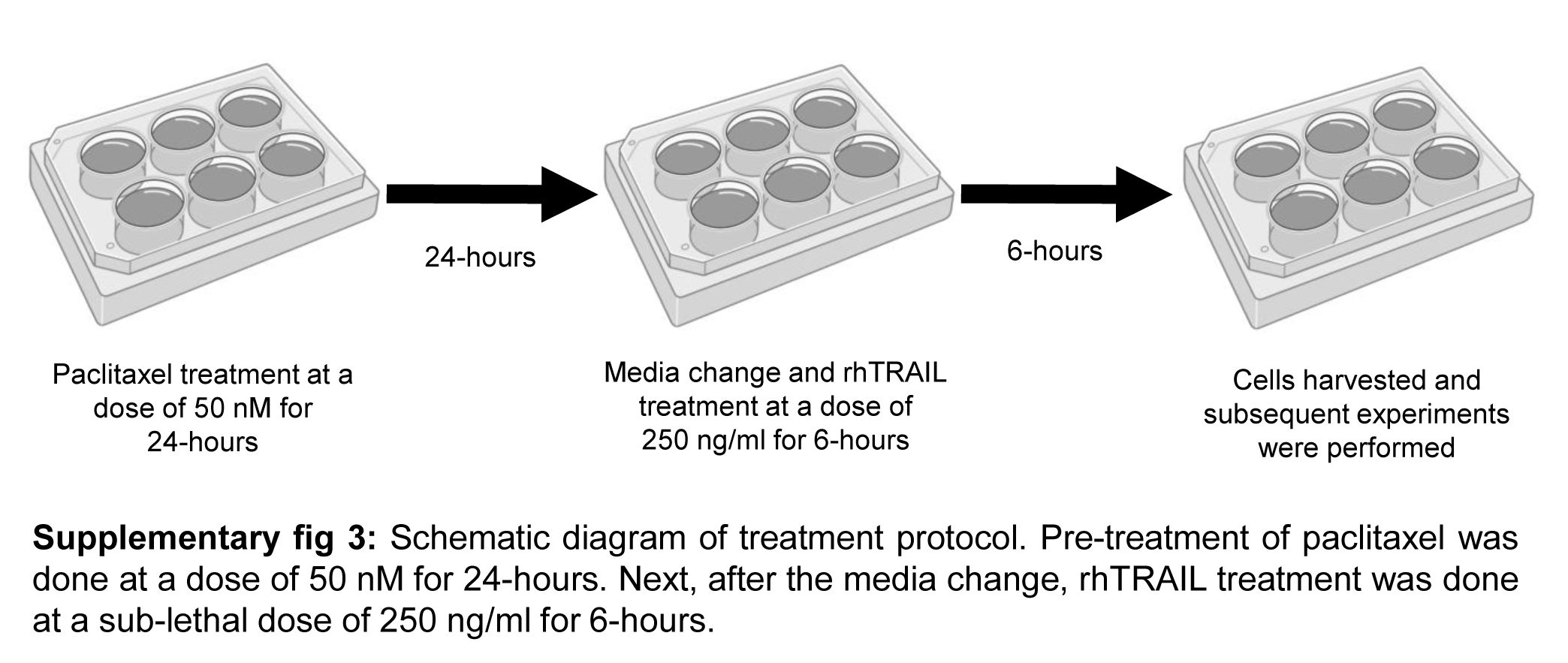
